# Texture-Analysis Classification of Intraductal Carcinoma of the Prostate Using Label-Free Multimodal Nonlinear Optical Imaging

**DOI:** 10.1101/2024.12.20.629794

**Authors:** Justin R. Gagnon, Christian Harry Allen, Mame-Kany Diop, Frédérick Dallaire, Frédéric Leblond, Dominique Trudel, Sangeeta Murugkar

## Abstract

Intraductal carcinoma of the prostate (IDC-P) is a very aggressive histopathological subtype of prostate cancer (PCa) for which no accurate biomarkers exist. In our work, we apply a multimodal nonlinear optical imaging approach that uses second-harmonic generation (SHG) and stimulated Raman scattering (SRS) imaging to distinguish IDC-P from regular PCa and benign prostate tissue. Images from each tissue type were classified using support vector machine (SVM). The technique classified the images from each region based on first-order statistics and texture-based second-order statistics derived from the gray-level co-occurrence matrix (GLCM) of the images. Our results demonstrate that SVM models trained on either SHG or SRS images accurately classify IDC-P as well as high-grade PCa, low-grade PCa, and benign tissue with a mean classification accuracy of over 89%. Furthermore, a classification model combining both SHG and SRS imaging modalities can accurately classify all tissue types with a mean classification accuracy of 98%.

## I. INTRODUCTION

Prostate cancer (PCa) accounts for more than one-fifth of diagnosed cancers in males and is the third-largest cause of cancer death in Canadian men [1]. A particularly aggressive histopathological subtype of PCa is intraductal carcinoma of the prostate (IDC-P). Due to the poor clinical outcomes of IDC-P in comparison to less aggressive subtypes of regular PCa, it is urgently necessary to provide pathologists with accurate biomarkers capable of discriminating IDC-P from regular PCa.

Nonlinear optical (NLO) imaging modalities have risen in prominence as diagnostic tools due to the non-ionizing nature of photon interactions and their ability to provide detailed biomolecular information at microscopic scales and high speed without the requirement for tissue staining dyes [2, 3]. In particular, stimulated Raman scattering (SRS) microscopy is a proven NLO chemical imaging technique [4, 5] that utilizes infrared light to excite and produce images from the different vibration modes of chemical bonds in biological samples. It has seen a multitude of applications including virtual histology [6, 7], in vitro [8], and in vivo [9] biological imaging. Second-harmonic generation (SHG) microscopy is another powerful NLO imaging technique that specializes in label-free collagen imaging [10] due to its specificity in detecting only noncentrosymmetric structures and has been used to diagnose diseases such as cancer which alter collagen organization [11–13].

SRS has previously been used to evaluate cholesteryl ester content to differentiate low-grade and high-grade PCa [14]. It has also been used to analyze cellular lipids and proteins in the carbon-hydrogen (CH) stretching “high-wavenumber” region of the Raman spectrum for PCa Gleason scoring [15], a scoring system used to quantify the grade of PCa. SHG has similarly been applied for PCa grading based on collagen fiber architecture [16–18] using techniques that evaluate the degree of collagen fiber orientation and alignment. However, neither of these imaging modalities have previously been used to identify IDC-P. Furthermore, none of these studies considered a multimodal approach combining SHG and SRS modalities to exploit their synergistic diagnostic potential. Here, we applied a multimodal approach that investigated the use of SRS as well as SHG imaging to distinguish IDC-P from high-grade PCa (HGC, Gleason grade 4 or 5), low-grade PCa (LGC, Gleason grade 3) and benign prostate tissue. Based on Raman biomarker band information identified earlier using Raman spectroscopy [19] we collected SRS images at the 1450 cm^−1^ and 1668 cm^−1^ Raman shifts in the “fingerprint” region of the Raman spectrum, followed by SHG images of the same field of view (FOV) from 50 PCa tissue biopsies. Texture analysis [20], a method that quantifies textural features of images based on gray-level differences, has previously been applied to the analysis of collagen and tissue organization for detecting cancer alongside a variety of NLO imaging modalities including SHG, coherent anti-Stokes Raman scattering, and two-photon excited fluorescence [21–23]. The goal of our study was to determine whether texture analysis-based features of PCa tissue extracted from SHG images and SRS images at the above Raman bands can be useful for the high-throughput identification of IDC-P.

By combining texture analysis, a technique that has never before been applied to SRS images, and support vector machines (SVM)-based classification [24], we show that both SHG and SRS microscopy imaging, when taken individually, can discriminate the four aforementioned types of tissue (IDC-P, HGC, LGC, benign prostate). Our work also demonstrates that higher classification accuracies can be achieved when the information from both modalities is combined into a single SVM model. These results lay the groundwork for future clinical applications of nonlinear optical imaging for diagnosing IDC-P and PCa.

## I. MATERIALS AND METHODS

### A. Human Tissue Samples

This study was approved by the Centre hospitalier de l’Université de Montréal (CHUM) ethics review board (reference number: 15.107). A cohort of 50 patients diagnosed with PCa who underwent first-line radical prostatectomy and participated in the Canadian Prostate Cancer Genome Project (CPC-GENE) were included [25]. Formalin-fixed paraffin-embedded (FFPE) radical prostatectomy specimens from these patients were used to construct tissue microarrays (TMAs). Two consecutive 4 µm tissue sections from each TMA block were cut and mounted onto glass slides (an H&E image from one such glass slide is shown in Fig. S1 in Supplementary Information S1). The first sections underwent immunohistochemistry (IHC) staining to detect α-methylacyl-CoA racemase (AMACR)/p63/34BE12, followed by hematoxylin and eosin (H&E) counterstaining [26]. The second sections were unstained and used for SHG and SRS imaging. IHC-H&E slides were scanned using a whole-slide scanner (Nanozoomer, Hamamatsu). Images of selected regions were classified as IDC-P, LGC, HGC, or benign. IDC-P was identified according to Guo and Epstein established criteria and confirmed by IHC staining [27]. Images with invasive carcinoma were classified and annotated by a pathologist based on the highest Gleason grade in the image: LGC if they contained only Gleason grade 3 architecture, or HGC if any Gleason grade 4 or 5 patterns were present. Finally, images were classified as benign if they contained only benign glands. In total, IDC-P images were obtained from 18 patients, HGC from 28 patients, LGC from 15 patients, and benign images from 5 patients.

### B. Multimodal NLO Imaging

The multimodal NLO images analyzed in this study were collected using a custom-built multimodal SRS microscopy platform. In this setup, an optical parametric oscillator system (Insight DS+, Spectra-Physics) provides a fixed laser output (“Stokes”; 1040 nm, *∼* 200 fs) and a tunable laser output (“pump”; 680-1300 nm, *∼* 130 fs) at a repetition rate of 80 MHz to target the Raman band of interest. As described in detail elsewhere (See [28, 29]), the spectral focusing-based SRS imaging platform employs a novel adjustable-dispersion glass blocks technology. The high refractive index (TIH53) glass blocks chirp the pump and Stokes pulses to 2.50 ps and 1.32 ps respectively, and the temporal delay between both excitation laser pulses is changed so that the instantaneous frequency difference matches that of the target Raman band. A 0.75NA 20X air objective (U2B825, Olympus) was used to image the tissue samples on regular microscope slides in transmission mode. The corresponding change in intensity of the pump beam due to stimulated Raman loss was measured as the SRS signal using lock-in detection. ScanImage (Version 5.6, Vidrio Technologies) was used to control laser scanning [30]. The SRS microscopy setup was modified to also collect SHG images by adding a longpass dichroic mirror with a cut-on wavelength of 450 nm (ZT1040dmbp, Thorlabs) to the output path that directed the second-harmonic light to another PMT (C14455-1560, Hamamatsu) as illustrated in Fig. 1(b). Following the collection of the SRS images in the region of interest, the SHG images of the same FOV were collected by tuning the chirped pump laser to a wavelength of 830 nm. A time-averaged pump beam power of 65 mW at the sample was used to collect both the SHG and SRS images, while the SRS images also used a Stokes beam with a time-averaged power of 45 mW. The NLO imaging location was selected based on a side-by-side comparison of real-time 1450 cm^−1^ SRS images and the H&E image of the same sample core annotated by the pathologist (e.g. Fig. 1(a)) for regions of IDC-P, HGC, LGC, and benign prostate tissue. Each imaged location was additionally verified by a pathologist post-collection to ensure the tissue label assigned to the locations was accurate. The SRS images were collected in the fingerprint region at the Raman bands of 1450 cm^−1^ and 1668 cm^−1^, corresponding to pump wavelengths of 904 nm and 886 nm. The 1450 cm^−1^ band represents the CH_2_ deformation vibrational mode with contributions from proteins and lipids while the 1668 cm^−1^ band represents the C=O stretch of the amide I vibrational mode with contributions from proteins [31]. These two Raman bands were chosen due to their known importance in classifying IDC-P and invasive PCa tissue [19] in addition to their prominent peak heights providing a strong signal-to-noise ratio (SNR) in the SRS images relative to other Raman bands in the fingerprint region. The vibrationally resonant Raman signal in SRS images is typically accompanied by a non-vibrationally resonant background signal which arises due to optical effects such as cross-phase modulation, two-photon absorption, and thermal lensing [32]. To suppress the background contributions, off-Raman resonance images were also collected in the valleys between Raman peaks based on Raman spectroscopy data in the prostate tissue [19] that represented only contributions from background effects. These off-Raman resonance images were collected at the Raman shifts of 1521 cm^−1^ (pump wavelength of 898 nm) and 1723 cm^−1^ (pump wavelength of 882 nm) corresponding to the on-Raman resonance images at 1450 cm^−1^ and 1668 cm^−1^ respectively. Images representing the vibrationally resonant Raman contribution were then obtained by subtracting the off-Raman resonance background image from the on-Raman resonance image. All SHG and SRS images were constructed by averaging 10 successively imaged frames at a resolution of 512×512. A pixel dwell time of 20 µs was used with a corresponding frame acquisition time of 5.2 s. The time required to acquire all SRS and SHG frames (in the following order: 1450 cm^−1^ on-resonance + off-resonance, 1668 cm^−1^ on-resonance + off-resonance, SHG) for one imaging location was approximately 6.5 minutes. Representative SHG and SRS images from the same 210 µm *×* 210 µm FOV in a prostate tissue core biopsy are shown in Fig. 1(c,d,e) and Fig. 2. In total, data was collected from 201 FOV regions among the different cores.

**FIG. 1.**
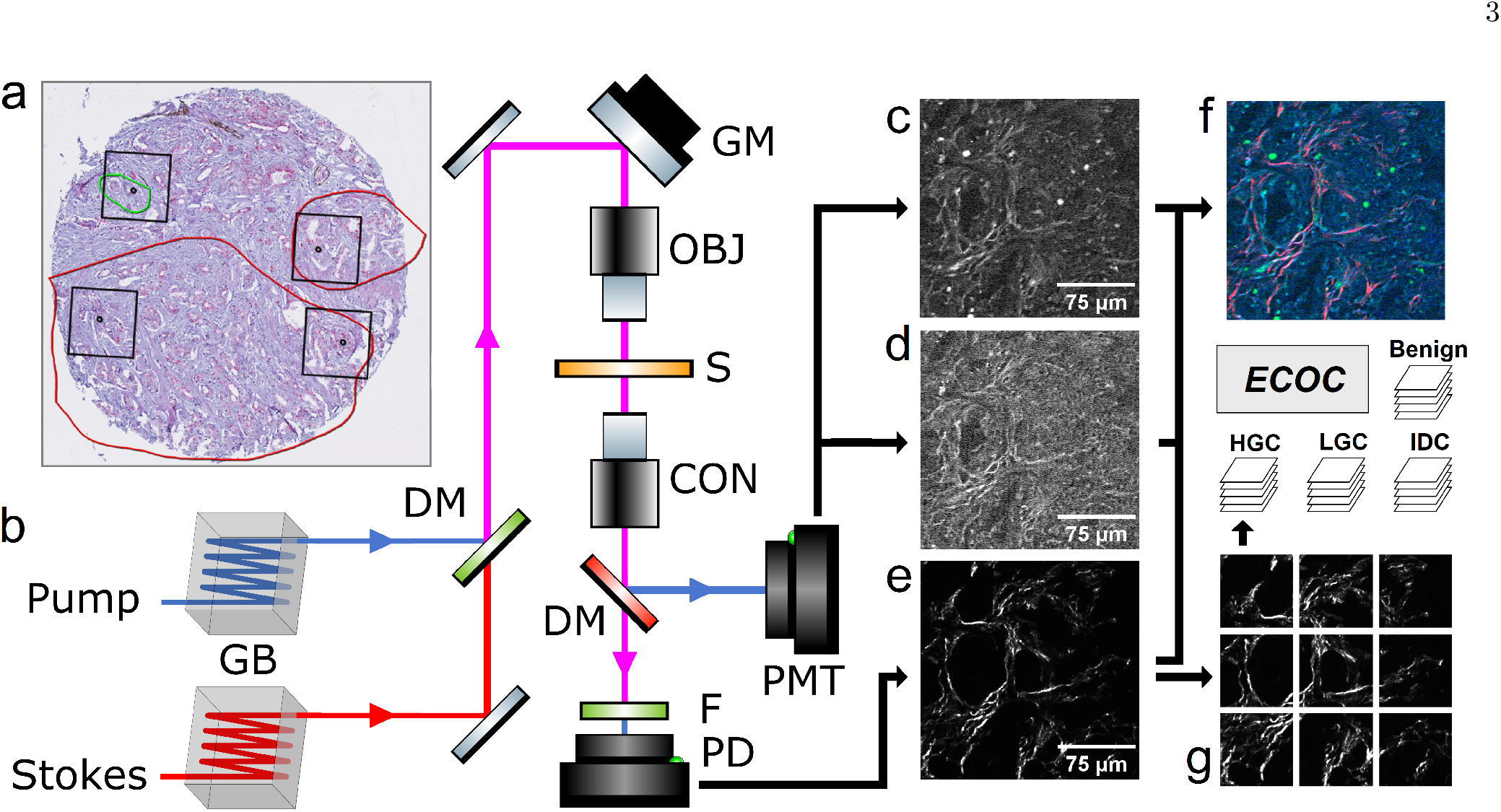
Experimental setup schematic and workflow. a) A representative annotated H&E image of a biopsied prostate tissue core with tissue regions annotated by a pathologist. HGC regions are shown in red and LGC regions in green, with the remainder being benign. Images were collected in the four regions demarcated by black squares on the core. In b) the experimental setup, the pump and Stokes pulses are chirped by glass blocks (GB), combined at a dichroic mirror (DM), and scanned through an objective (OBJ) onto the sample (S) using galvanometer mirrors (GM). The light is then collected using a condenser (CON). Another dichroic mirror redirects the SHG light to the PMT, and the remaining light is collected in the SRS path after filtering the 1040 nm light and focusing it on the photodetector. This process yields c) 1450 cm^−1^ SRS images and d) 1668 cm^−1^ SRS images after background subtraction, and e) SHG images. f) A co-registered image where SHG is shown in red, 1450 cm^−1^ SRS is shown in green, and 1668 cm^−1^ SRS is shown in blue. All the different multimodal NLO images (FOVs) are split into 3 × 3 subimages as shown in g) to train and test an SVM (ECOC) model.

**FIG. 2.**
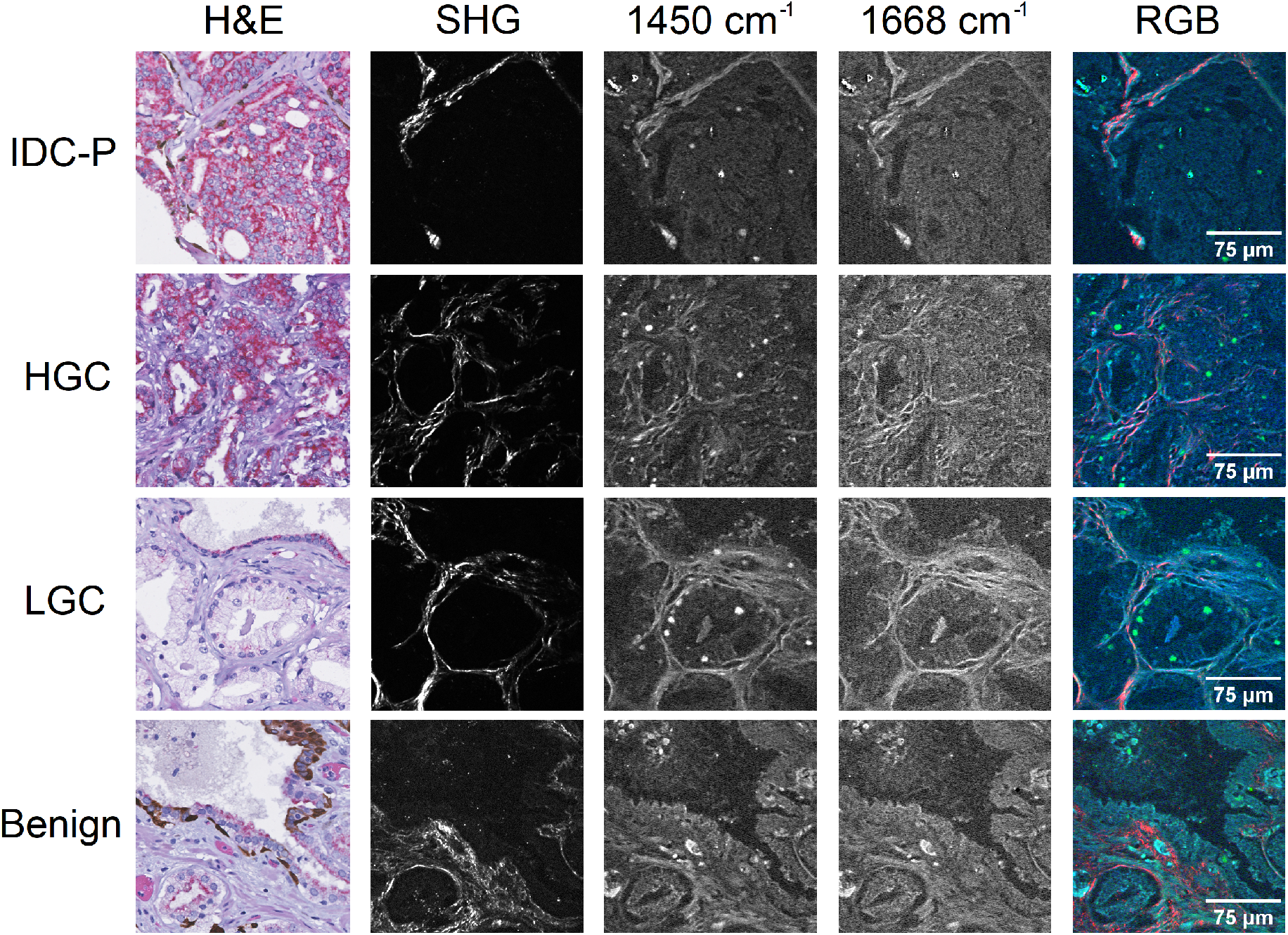
Representative images collected for different types of prostate tissue: IDC-P, HGC, LGC, and benign. The first column shows H&E/immunohistochemistry images for each tissue, and the three following columns show images of the same region acquired with SHG, with SRS at 1450 cm^−1^, and with SRS at 1668 cm^−1^. The final right-most column shows co-registered images where SHG is shown in red, 1450 cm^−1^ SRS is shown in green, and 1668 cm^−1^ SRS is shown in blue.

### C. Texture Analysis

To differentiate each different type of prostate tissue, the images were quantified using texture analysis based on first- and second-order statistics [24]. First-order statistics are directly derived from the gray-level pixel intensities of the image and do not consider the positioning of the pixels. In this study, the values of mean, variance, skewness, and kurtosis are used as first-order statistics. Second-order statistics are instead derived from the spatial relations between the different pixels. Second-order statistics are therefore a method of quantifying the textures of the image, extracting information beyond the gray-level pixel intensity information used in first-order statistics. The spatial relations between the pixels are described by the gray-level co-occurrence matrix (GLCM), which quantifies how often co-occurring pixel values appear with specific intensities at a given pixel offset [24]. For each image, GLCMs were calculated for four different one-pixel offsets (horizontal, vertical, diagonally up, diagonally down) after min-max normalizing the image to 128 gray levels. To ensure that aberrant bright pixels did not influence the normalization, the top 1% of pixels were all assigned a gray level of 128. The second-order statistical parameters of contrast, homogeneity, energy, correlation, and entropy were then calculated by performing operations on the GLCM; an in-depth description of these statistical quantities can be found in Supplementary Information S2. Because the statistical quantities depend on the offset of the GLCM, each quantity was calculated for each of the four GLCM matrices and then averaged to yield a singular statistic.

### D. Image Segmentation for Data Analysis and FOV Scoring

To increase the number of images to train and test on, all collected FOVs were split into a 3×3 grid, with a final count of 267 IDC-P subimages, 544 HGC subimages, 221 LGC subimages, and 46 benign tissue subimages. A few FOVs contained more than one tissue type, in which case the image was split into two distinct FOVs corresponding to the different tissues. Other FOVs overlapped causing some of their subimages to coincide, in which case additional copies of a given subimage were removed. Due to the collagen imaged using SHG microscopy not always filling the entire image, split SHG subimages found not to contain enough collagen were discarded through a manual review. Fig. S3 in Supplementary Information S3 provides examples of subimage selection for the three above cases. The final count of SHG subimages came to 124 IDC-P subimages, 423 HGC subimages, 201 LGC subimages, and 34 benign tissue subimages. To account for the discrepancy in the number of subimages for each tissue type, the subimages were assigned learner weights such that the sum of all subimages for each tissue type had the same weight.

### E. Multi-Class Discrimination of Subimages and FOVs Based on SVM Model Training

The SVM consisted of an error-correcting output code (ECOC) model in MATLAB R2024a. ECOC models enable multiclass learning by combining multiple binary SVM learners (one for each set of two classes) to construct a classifier. The ECOC model was trained using a polynomial kernel on the combination of 9 first- and second-order statistical parameters for each imaging modality (SHG and both SRS bands). The SVM model was trained to discriminate between the four basic types of prostate tissue: IDC-P, HGC, LGC, and benign tissue. The training and test sets were determined using a quasi-random splitting approach, where the subimages were split in a 2:1 ratio to serve as the training and test sets respectively. Here, quasi-random refers to the fact that prostate cores in the training set did not appear in the test set and vice versa for each tissue type. This constraint eliminated the intra-core bias that would arise from having subimages from the same core in both sets. When performing the split, the subimages corresponding to each tissue were split 2:1 to ensure an equal representation of tissue types in the training and test sets. The quasi-random split was performed 10 times and the accuracy results from scoring the test set in each iteration were averaged to yield the final results. The accuracy is reported as a confusion matrix showing what percentage of subimages of a given tissue type (True class) are classified as a certain tissue type (Predicted class).

### F. Multi-Class Discrimination of FOVs Based on SVM Model Training

An alternative approach to taking the classification accuracy of the subimages is to determine how the FOVs (the original images before being split into 3×3) are classified by the model. This is done by finding the majority consensus (mode value) of the subimages composing each FOV. The approach of classifying the FOV, as opposed to each subimage individually, allows the SVM model to provide a definitive diagnosis for an imaged area. The results for the FOV majority-consensus classification accuracy are reported as a separate confusion matrix. In the event of a tie between two tissue types, the FOV was classified as the most aggressive tissue (IDC-P *>* HGC *>* LGC *>* benign tissue). The approach of FOV classification furthermore provides a convenient method of implementing a multimodal approach to image classification. The multimodal model returned the confusion matrix obtained from the aggregate majority consensus of both the SHG and SRS subimages for a particular FOV.

### G. Feature Selection

It sometimes occurs that certain statistical quantities do not end up having discriminatory power and are not useful for distinguishing different tissue types. When this is the case, these quantities only add noise to the model, decreasing its classification accuracy or potentially leading to over-fitting. To account for this possibility, one model was trained using all parameters, and then models were also trained where all parameters except one were present. When the exclusion of a parameter was found to increase the classification accuracy relative to the initial model, this parameter was excluded from the final model. The conditions under which these one-parameter-excluded models were trained were identical to the main model, that is to say, a 2:1 quasi-random split where the final results were averaged over 10 iterations.

## II. RESULTS

### A. SHG and SRS Images of Prostate Tissue

Representative images for IDC-P, HGC, LGC, and benign prostate tissue imaged in the experiment are shown in Fig. 2. When comparing to the H&E images, it can be seen that the SHG images depict the collagen architecture of the imaged regions. The 1450 cm^−1^ and 1668 cm^−1^ SRS images are visually more similar to the H&E images since all components of the tissue contain the molecular signatures inherent to both SRS bands. When comparing both types of SRS images against one another, the 1450 cm^−1^ SRS images have higher SNR with stronger contributions from the stroma, and additional bright spots originating from lipofuscin granules [14]. In contrast, 1668 cm^−1^ SRS images show increased structural detail from protein contributions surrounding the stroma. Fig. 2 also presents co-registered images which show the relative signal intensities of all three NLO channels after contrast enhancement.

### B. Testing the Discriminatory Power of GLCM Statistics

Before training an SVM model based on the first- and second-order parameters, it is important to verify that statistical differences in those parameters are present between the different types of prostate tissue. This was done by using the Kruskal-Wallis test, a non-parametric ranked sum test that returns the *p*-value that different statistics come from the same distribution. The GLCM calculations and Kruskal-Wallis test were performed in MATLAB R2024a (Mathworks) for the SHG, 1450 cm^−1^ SRS, and 1668 cm^−1^ SRS subimages. The test evaluated, for each pairing of tissues (for example, IDC-P versus HGC) and for each statistic derived from the subimages, the *p*-value that the statistics did not originate from a single distribution. The results are provided in Tables S1, S2, and S3 respectively in Supplementary Information S4, and generally indicate good discriminatory power between the different tissue types due to multiple statistics having statistically significant *p*-values (*p <* 0.05) for most tissue pairings. More statistically significant *p*-values were present in the tissue pairings for SHG subimages (69% of statistics) than in the 1450 cm^−1^ SRS subimages (54.8% of statistics) and 1668 cm^−1^ SRS subimages (57.1% of statistics).

### C. Feature Selection Results

The comprehensive data showing the effects of the feature selection process are shown in Table I. For the SVM model trained on SHG subimages, omitting the parameters of variance, skewness, kurtosis, or energy led to the one-parameter-removed model exceeding the classification accuracy of the baseline model, so these parameters were removed from the final SVM model. In the SVM model trained on 1450 cm^−1^ SRS subimages, all one-parameter-removed models exhibited lower overall classification accuracies, so all parameters were kept for the final SVM model. In the SVM model trained on 1668 cm^−1^ SRS subimages, omitting the parameters of mean and kurtosis increased the overall classification accuracy. Omitting the noisy parameters had the effect of increasing the overall classification accuracy of the SVM model trained on SHG subimages from 92.6% to 94.2%, and increasing the overall classification accuracy of the SVM model trained on 1668 cm^−1^ SRS subimages from 85.4% to 89.1%, a sizeable increase in both cases.

**TABLE I.**
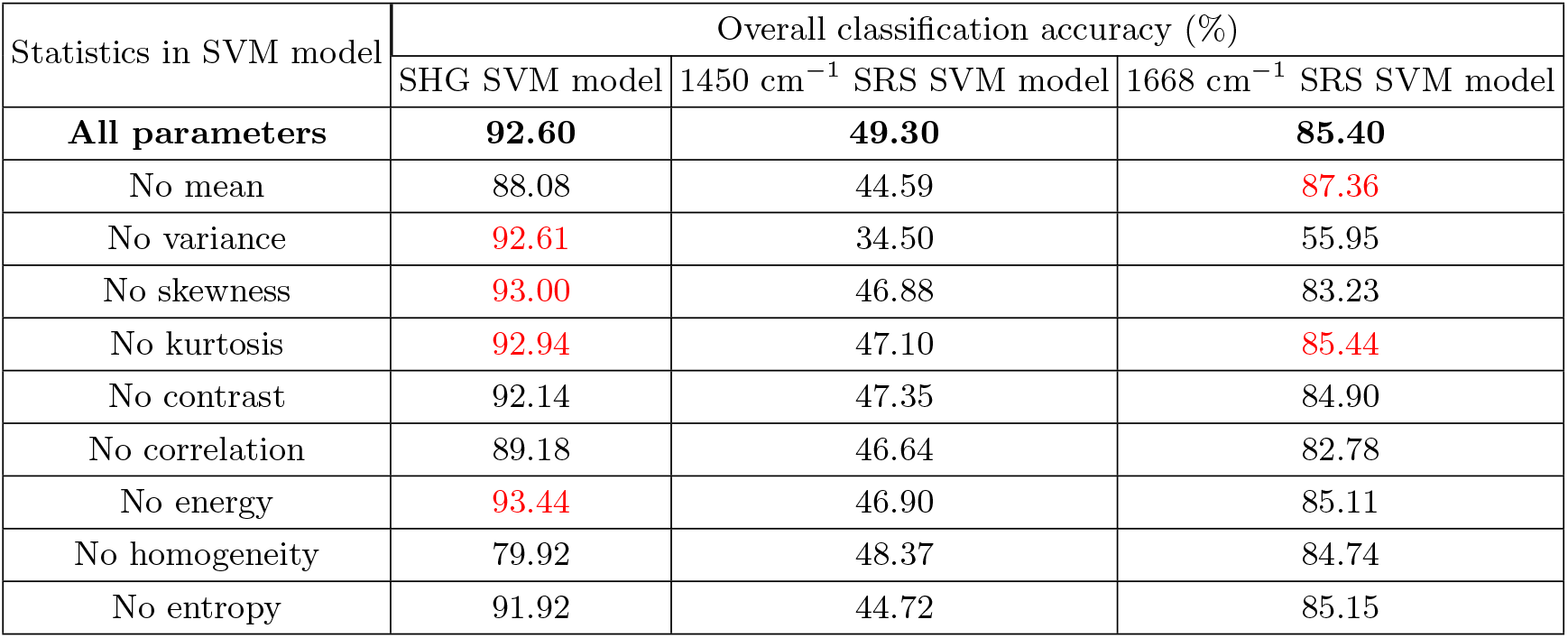
Results for feature selection. Mean classification accuracy of the central diagonal of the confusion matrix representing subimage accuracy as a function of which parameters are trained on. Red quantities are classification accuracies that are higher than when using all parameters (shown in bold), indicating that omitting that statistic increased the classification accuracy of the model.

| Statistics in SVM model | Overall classification accuracy (%) |  |  |
| --- | --- | --- | --- |
|  | SHG SVM model | 1450 cm<sup>-1</sup> SRS SVM model | 1668 cm<sup>-1</sup> SRS SVM model |
| **All parameters** | **92.60** | **49.30** | **85.40** |
| No mean | 88.08 | 44.59 | **87.36** |
| No variance | **92.61** | 34.50 | 55.95 |
| No skewness | **93.00** | 46.88 | 83.23 |
| No kurtosis | **92.94** | 47.10 | **85.44** |
| No contrast | 92.14 | 47.35 | 84.90 |
| No correlation | 89.18 | 46.64 | 82.78 |
| No energy | **93.44** | 46.90 | 85.11 |
| No homogeneity | 79.92 | 48.37 | 84.74 |
| No entropy | 91.92 | 44.72 | 85.15 |

### D. Multi-Class Discrimination of Subimages in the Three Channels

The confusion matrices describing the classification accuracies of the subimages for the SVM models trained on SHG, 1450 cm^−1^ SRS, and 1668 cm^−1^ SRS subimages are shown in Fig. 3(a,b,c). The SVM model trained on SHG subimages reports high classification accuracies for subimages of each tissue type. Notably, the classification accuracies of HGC, LGC, and benign tissues all exceed 94%. However, there is a notable weakness in this model for detecting IDC-P, with 11.7% of IDC-P subimages being misclassified as HGC. This weakness can be linked to the Kruskal-Wallis test result for SHG subimages (Table S1), which returned high *p*-values for all statistics when comparing IDC-P and HGC. The classification accuracies for the SVM model trained on 1450 cm^−1^ SRS subimages are substantially lower than those of the SVM trained on SHG subimages, with a mean classification accuracy of only 49.3%. In contrast, the SVM model trained on 1668 cm^−1^ SRS subimages achieves classification accuracies similar to those of the SVM model trained on SHG subimages, with classification accuracies exceeding 92% for IDC-P and HGC tissues. However, this SVM model misclassifies 16.7% of benign tissue subimages as IDC-P and misclassifies 14.7% of LGC tissue as either HGC or IDC-P. This weakness can be linked to the Kruskal-Wallis result for the 1668 cm^−1^ SRS subimages, which showed IDC-P and benign tissues had only one statistically significant statistic.

**FIG. 3.**
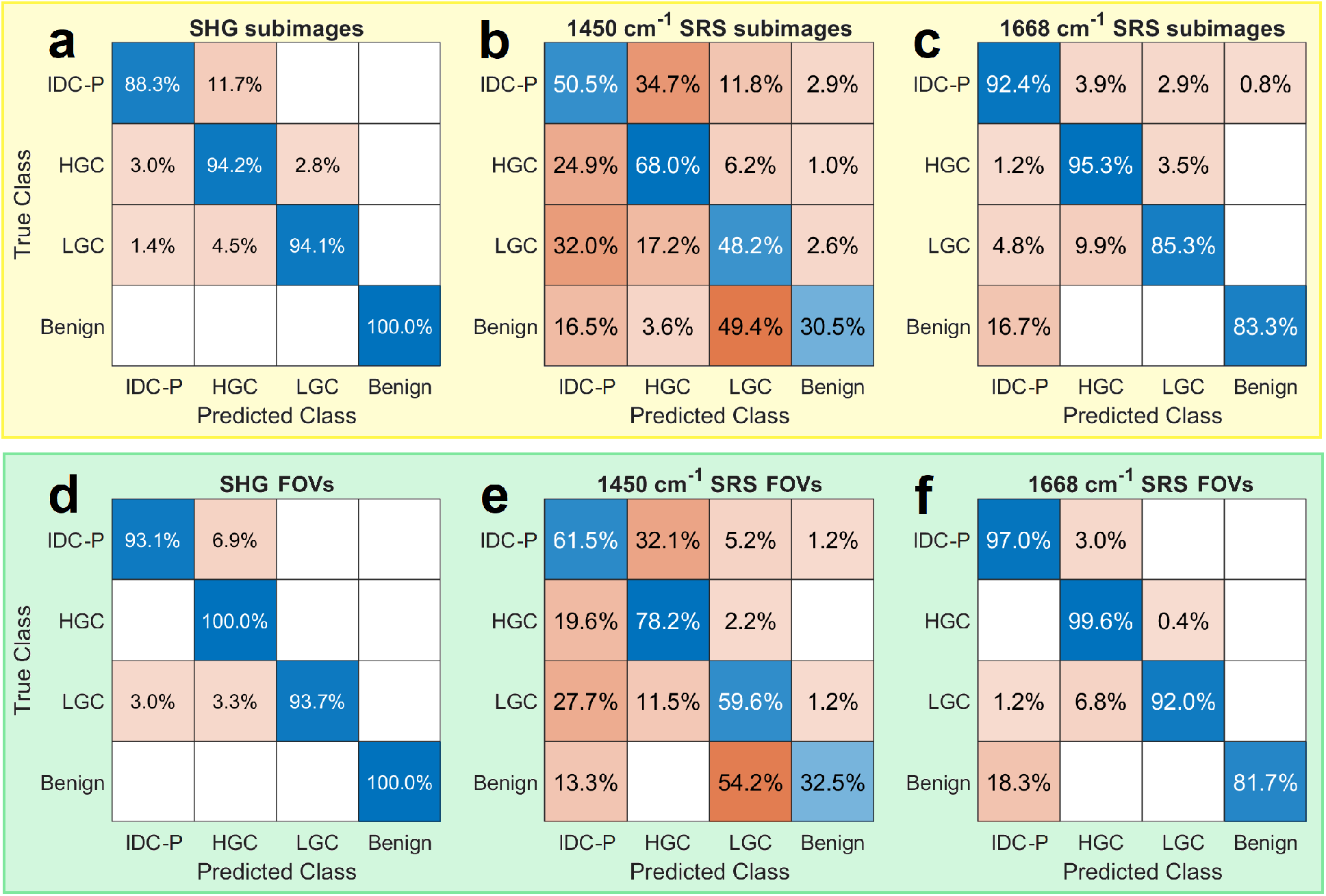
Confusion matrices for classification of a) SHG, b) 1450 cm^−1^ SRS, and c) 1668 cm^−1^ SRS subimages. The confusion matrices obtained when incorporating an FOV majority-consensus model are shown for d) SHG, e) 1450 cm^−1^ SRS, and f) 1668 cm^−1^ SRS FOVs. The average classification accuracies of the central diagonal for each confusion matrix are 94.2% and 96.7% for the SHG models (subimage and FOV), 49.3% and 58% for the 1450 cm^−1^ SRS models, and 89.1% and 92.6% for the 1668 cm^−1^ SRS models.

### E. Multi-Class Discrimination of FOVs Based on Majority Consensus

Incorporating an FOV majority-consensus model generally increases the classification accuracy for each tissue type (Fig. 3(d,e,f)). Most notably, the mean classification accuracy increased for each imaging modality, with the mean classification accuracy increasing from 94.2% to 96.7% for the SHG model, from 49.3% to 58% for the 1450 cm^−1^ SRS model, and from 89.1% to 92.6% for the 1668 cm^−1^ SRS model. In the case of the SVM model trained on SHG subimages, the number of IDC-P subimages classified as HGC is reduced from 11.7% to 6.9%, at the cost of a slightly higher fraction (5.9% to 6.3%) of misclassified LGC subimages. In the case of the SVM model trained on 1450 cm^−1^ SRS subimages, the majority-consensus model increases the classification accuracies of all categories, but still only achieves a low mean classification accuracy of 58%. For the SVM model trained on 1668 cm^−1^ SRS subimages, the classification accuracy for all cancerous tissues increases to above 92%, but the classification accuracy for benign tissue sees a slight decrease to 81.7%.

### F. Multimodal SHG and SRS-Based Multi-Class Discrimination of FOVs

Due to the low classification accuracy of the SVM model trained on 1450 cm^−1^ SRS subimages, the multimodal SHG + SRS model was implemented by using the SHG and 1668 cm^−1^ SRS subimages. The multimodal model therefore classified the FOVs by evaluating the majority-consensus of the combined SHG and 1668 cm^−1^ SRS subimages taken in each particular FOV. The confusion matrix for the multimodal multi-class discrimination SVM model is shown in Fig. 4. It has nearly perfect accuracy, correctly classifying all tissues at accuracies exceeding 96%, and with an average classification accuracy of 98%. Notably, the multimodal SVM model has 2.6 times fewer IDC-P misclassifications than the SHG majority-consensus model (Fig. 3(d)) with an increase in classification accuracy from 93.1% to 97.4%. Moreover, it shows 8.7 times fewer misclassifications for benign tissue than the 1668 cm^−1^ SRS majority-consensus model (Fig. 3(f)) with an increase in classification accuracy from 81.7% to 97.9%. These increases in classification accuracy indicate that the multimodal SVM model mitigates the classification weaknesses of SVM models using only one modality.

**FIG. 4.**
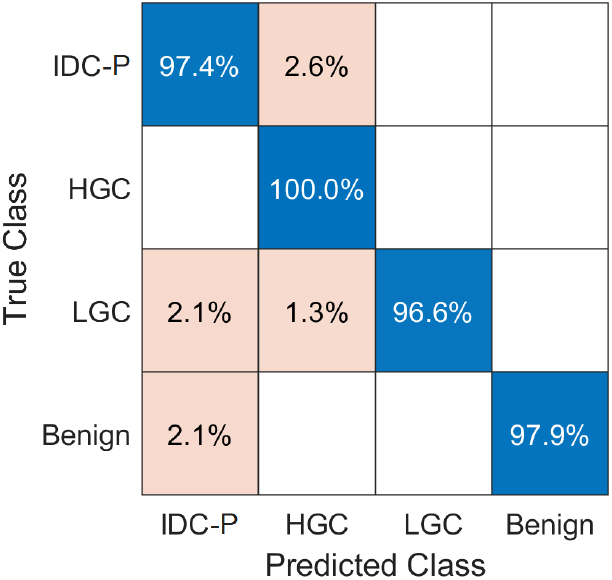
Confusion matrix for multimodal FOV majority-consensus classification of the SHG and 1668 cm^−1^ SRS subimages. The average classification accuracy of the central diagonal of the confusion matrix is 98%.

## III. DISCUSSION

In this work, we developed a method to discriminate between subtypes of PCa tissue using an SVM model trained on texture analysis features extracted from SHG and SRS images at the 1450 cm^−1^ and 1668 cm^−1^ Raman bands. The first- and second-order statistics trained on by the SVM models showed a high proportion of low *p*-values when using the Kruskal-Wallis test, indicating the high discriminatory power of the statistics.

The SVM model trained on SHG subimages reported high classification accuracies for each tissue type (*>*88%) that were even further improved after implementing FOV majority-consensus classification (*>*93%). As seen in Fig. 3(a), using an FOV majority consensus improved the results by reducing the percentage of misclassifications of IDC-P as HGC from 11.7% to 6.9%. The high mean classification accuracies of 94.2% with subimages and 96.7% with FOV majority-consensus are consistent with the established potential of SHG for PCa grading [16–18]. However, in contrast to these prior studies, our SHG results establish the potential of SHG to identify IDC-P on top of achieving accurate grading for regular PCa, suggesting that changes in collagen architecture can be considered a reliable biomarker for IDC-P.

The SVM model trained on 1450 cm^−1^ SRS subimages demonstrated much lower classification accuracies when compared to the SVM models trained on SHG and 1668 cm^−1^ SRS subimages. The low classification accuracies disqualify the possibility of using SRS images at the 1450 cm^−1^ Raman band for PCa scoring and identification of IDC-P. However, the fact that the 1668 cm^−1^ Raman band was successful in contrast to the 1450 cm^−1^ band permits us to infer that the chemical bonds targeted by the 1668 cm^−1^ SRS band provide textural information that is useful for classifying cancerous and non-cancerous tissue that is not present in the 1450 cm^−1^ band. Fig. 2 shows the micromorphology in these two types of SRS images highlighting the different textural contributions of the two bands. In the 1668 cm^−1^ SRS image, the structure of the protein contributions surrounding the stroma is more detailed than in the 1450 cm^−1^ SRS image, despite the decreased SNR. Therefore, a probable reason for the success of the 1668 cm^−1^ SRS band is that the detailed cellular textures provide more discriminatory power. The success of the 1668 cm^−1^ band in contrast to the 1450 cm^−1^ band also highlights that only specific chemical components of the tissue, and therefore only images taken at certain Raman bands, can be used to discriminate cancerous and non-cancerous tissue types. During feature selection, the classification accuracy of both SVM models trained on SRS subimages sees a decline when the parameter of variance is omitted, with the accuracy of the 1450 cm^−1^ SRS SVM model decreasing by 14.8% and the accuracy of the 1668 cm^−1^ SRS SVM model decreasing by 29.4%. An explanation for the relative importance of the variance in these models is that the second-order parameters are correlated with one another: with only first-order parameters, the model achieves a classification accuracy of only 29.3%, as shown in Table S4 in Supplementary Information S5. Unlike the SHG images, which are composed purely of collagen, the SRS images are a combination of collagen and soft tissue. Since the collagen generally has a stronger signal than the soft tissue, a high variance is expected when both collagen and soft tissue are present while a low variance value is expected when there is only soft tissue. The variance may therefore be important for the SVM models trained on SRS subimages due to its ability to quantify the collagen to soft tissue ratio in the subimages.

While the individual SVM models trained on SHG subimages and 1668 cm^−1^ SRS subimages demonstrated a weakness at classifying IDC-P and benign tissue respectively, the multimodal SVM model incorporating the results of both SVM models does not demonstrate either shortcoming and provides a nearly perfect 98% average classification accuracy. The multimodal model therefore combines the classification strength of SHG for benign tissue and the classification strength of SRS for IDC-P. The lowest classification accuracy in the multimodal SVM model is 96.6% for LGC tissue, reflecting that the SHG image and 1668 cm^−1^ SRS image SVM models both had LGC as their second least accurate classification category. This lower accuracy may be attributable to LGC regions being in proximity to HGC. However, the LGC classification percentage of the multimodal SVM model still considerably exceeds those obtained by the SHG majority-consensus model (93.7%) and 1668 cm^−1^ SRS majority-consensus model (92.0%), demonstrating the considerable benefit of a multimodal approach.

The results exhibited in this paper involved imaging SRS bands in the fingerprint region (800–1800 cm^−1^) with the intention of building on a previous work that demonstrated that several Raman bands in the fingerprint region had the potential to discriminate IDC-P from cancerous and benign tissue [19]. However, Raman bands in the CH stretching high-wavenumber region (2800–3100 cm^−1^) are typically more convenient to image using SRS due to the much higher density of CH bonds giving rise to a stronger SRS signal and therefore higher SNR. Potential Raman bands of interest in the CH region include the lipid, overlapping protein and lipid, and DNA bands located at 2850 cm^−1^, 2926 cm^−1^, and 2970 cm^−1^ respectively. A convolutional neural network trained on SRS images at the lipid and mixed protein and lipid bands demonstrated a Gleason scoring accuracy of 84.4%, this suggests that these bands could also be useful for detecting IDC-P [15]. We anticipate that our approach based on textural analysis of SHG and SRS images when extended to the Raman bands in the high-wavenumber region will have the advantage of providing *>*95% classification accuracy of IDC-P at significantly reduced (at least 2x lower) image acquisition times due to the increased SNR. The set of images used in this experiment included a low proportion of benign tissue due to the reclassification of some tissue as cancerous by pathologists following data collection. While the current study provides compelling results showing that texture analysis is fully capable of classifying benign tissue from cancerous tissues, future studies will expand the assortment of benign images to further bolster this claim.

A conclusive diagnosis of IDC-P is difficult for pathologists due to subjective diagnostic criteria and a lack of available biomarkers [33]. The results we present in this paper exhibit that NLO imaging techniques based on SHG and SRS imaging can be combined with texture analysis and SVM classification to provide pathologists with a reliable biomarker of IDC-P. Notably, both SHG and SRS (1668 cm^−1^) imaging modalities individually demonstrated high classification accuracies, while nearly perfect classification accuracies were achieved by combining both modalities. The clear benefit obtained from using two imaging modalities to classify IDC-P further highlights the potential of multimodal SHG + SRS acquisitions in a clinical setting. Due to the use of 4 µm FFPE tissue sections and standard glass slides, our method is also compatible with conventional histopathological workflow [19]. Avenues for future research involve expanding these methods to a broader set of data for further validation and clinical applications such as an intraoperative handheld probe for the diagnosis of disease from prostate cores obtained in vivo.

## IV. ACKNOWLEDGEMENTS

We thank the University Health Network team for providing the TMAs, and Dr. Nazim Benzerdjeb MD who created the TMAs. We thank the molecular pathology core facility and Mirela Birlea of the CRCHUM for help in preparing prostate sections. Biobanking at the CRCHUM, CHUQc-UL, and MUHC was done in collaboration with the Réseau de Recherche sur le cancer of the Fonds de Recherche Québec - Santé (FRQS), which is affiliated with the Canadian Tumor Repository Network (CTRNet). The authors acknowledge the support of the Canadian Institutes of Health Research (CIHR) (funding reference number PJT-169164), and the Natural Sciences and Engineering Research Council (NSERC) of Canada (funding reference number RGPIN-2022-04897) (SM). Dr. Dominique Trudel receives salary support from the Fonds de Recherche du Québec, Santé (FRQS, Clinical Research Scholar, Senior). The CRCHUM also receives support from the FRQS.

