## Supplementary Information for "Texture-Analysis Classification of Intraductal Carcinoma of the Prostate Using Label-Free Multimodal Nonlinear Optical Imaging"

(Dated: January 9, 2025)

This document provides Supplementary Information for "Texture-Analysis Classification of Intraductal Carcinoma of the Prostate Using Label-Free Multimodal Nonlinear Optical Imaging". In Section S1, we provide a representative H&E image showing a slide of the patient biopsy cores. In Section S2, we provide background details on the calculation of the GLCM and its associated second-order parameters. In Section S3, we show three examples of cases occurring during image segmentation where data is discarded to decrease bias. In Section S4, we provide the p-values obtained when comparing each statistic between different tissue types when using the Kruskal-Wallis ranked sum test. In Section S5, we evaluate the relative importance of the first- and second-order parameters to the classification accuracies of the SVM models.

### S1. REPRESENTATIVE H&E IMAGE OF SLIDE WITH PATIENT BIOPSY CORES

The imaged prostate tissue cores were mounted onto a total of 5 glass slides. A representative slide is shown in Fig. S1. Each slide contained a 6 x 9 array of prostate cores, except for the fifth slide, which had a 7 x 12 array. Not all cores were included in the study, as images were only collected from cores with regions of interest.

### S2. DERIVATION OF GLCM AND CALCULATION OF SECOND-ORDER PARAMETERS

#### S2.1. Gray-Level Co-occurrence Matrix

The gray-level co-occurrence matrix (GLCM) quantifies texture in an image by considering the relations between what are known as *reference* and *neighbor* pixels. An example image for which a GLCM is calculated is shown in Fig. S2(a)

---

[*] D. Trudel and S. Murugkar contributed equally to this work

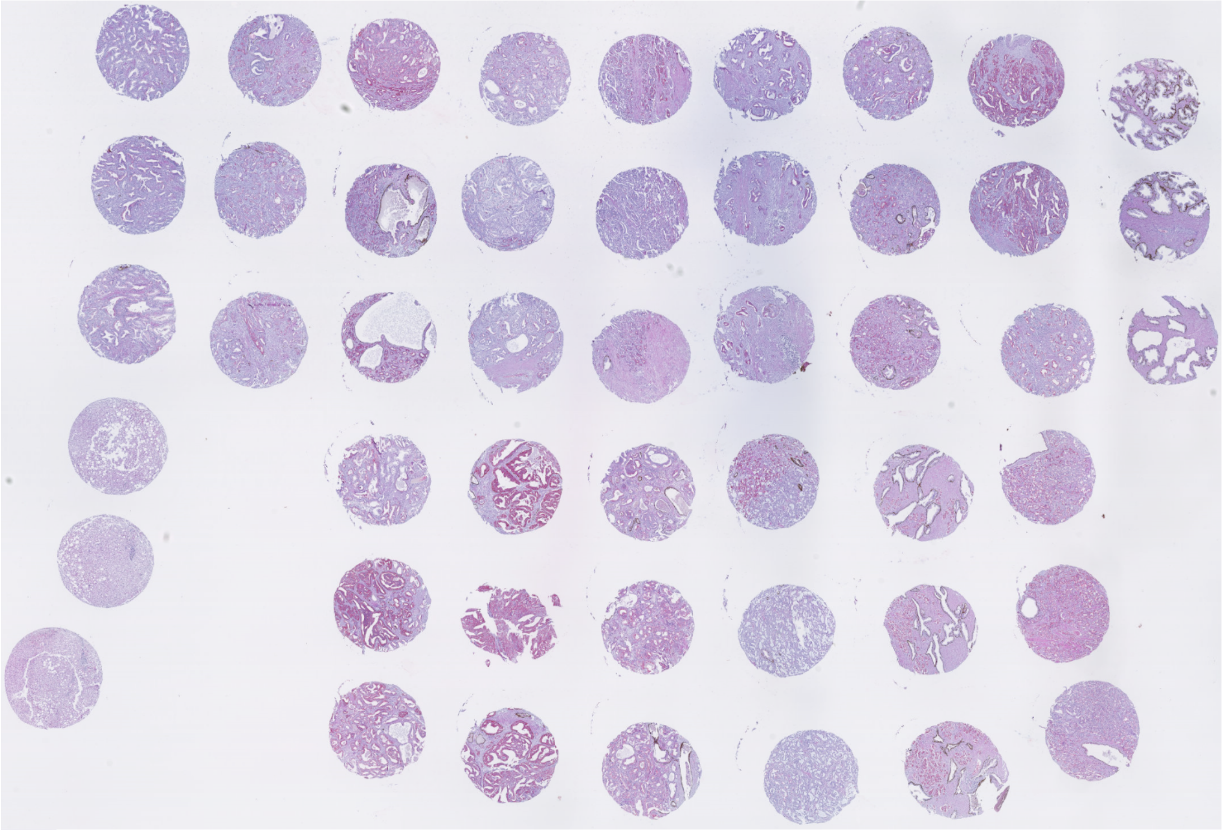

FIG. S1 Representative H&E image of a slide containing the patient biopsy cores.

with gray-level values shown in Fig. S2(b). The resulting GLCM in Fig. S2(c) quantifies the spatial relations between those gray values for a rightward offset between the pixels (that is to say, the neighbor pixel value is one pixel to the right of the reference pixel). For example, the entry in the GLCM corresponding to (2,2) has a value of 3 because a gray value of 2 occurs to the right of another gray value of 2 three times in the image.

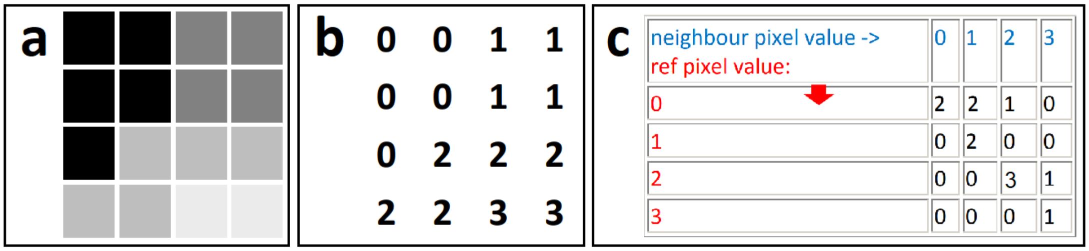


FIG. S2 A representative image with 4 gray levels is shown in a) and quantified in b). The resulting GLCM in c) is calculated by counting how many times a neighbor pixel with a certain gray value appears next to a reference pixel of a given gray value. Adapted from (Hall-Beyer, 2017).

Before calculating the texture parameters, the GLCM is normalized such that the sum of all its values is unity and that each element therefore represents the joint probability value between two pixels (Hall-Beyer, 2017). One feature of note of the GLCM is that its matrix dimensions are equal to the number of gray levels in the image. In this work, the prostate images are normalized to 128 gray levels before calculating the GLCM to minimize the number of zero values in the GLCM and increase its validity as an approximation of the joint probability value (Hall-Beyer, 2017). Another important feature is that elements on the central diagonal indicate that pixels have an identical value to their neighbor, while elements further from the central diagonal indicate pixels that differ progressively more in value from one another. This fact will form the basis for some of the texture parameters.

In this work, the GLCMs are made symmetrical before performing texture calculations. A GLCM matrix can be made symmetrical by adding it to its transpose. For the example image in Fig. S2, the transpose GLCM matrix would be calculated by considering neighbor pixels to the left of the reference pixel instead of to the right. The sum of both matrices is symmetrical about the diagonal. This form of the GLCM is more accurate to use in texture analysis since all pixels are considered as a reference and neighbor pixel in the calculation (for example, in the GLCM in Fig. S2c, the pixels on the right are never considered as reference pixels since they have no rightward neighbor). Furthermore, the symmetrical matrix in this case would be representative of all horizontal textures.

#### S2.2. Second-Order Texture Parameters

Second-order texture parameters can be calculated from the GLCM that provide quantities to evaluate the texture in the original image. To ensure that pixel relationships are evaluated omnidirectionally, these quantities are calculated for GLCMs based on horizontal, vertical, diagonally up, and diagonally down one-pixel offsets, and are then averaged together to yield the final quantity.

##### 1. Contrast

The quantity of contrast is given by Equation S1, where $N$ is the number of gray levels, and $P_{i,j}$ is the value of the GLCM element in column $i$ and row $j$. Due to the factor of $(i-j)^2$, elements further from the central diagonal of the GLCM are weighed more heavily in the calculation, and the contrast value being larger is therefore an indication of a larger disparity between reference and neighbor pixel values in the GLCM.

$$\sum_{i,j=0}^{N-1} P_{i,j}(i-j)^2 \tag{S1}$$

##### 2. Correlation

The quantity of correlation is given by Equation S2, where $\mu_i$ and ${\sigma_i}^2$ are the mean and variance of the GLCM respectively. The correlation parameter measures the dependency of gray levels on those of neighboring pixels and increases when there is a high predictability of pixel relationships.

$$\sum_{i,j=0}^{N-1} P_{i,j}\left(\frac{[i-\mu_i][j-\mu_j]}{{\sigma_i}^2{\sigma_j}^2}\right), \qquad \mu_i = \sum_{i,j=0}^{N-1} iP_{i,j}, \qquad {\sigma_i}^2 = \sum_{i,j=0}^{N-1} P_{i,j}(i-\mu_i)^2 \tag{S2}$$

##### 3. Energy

The quantity of energy is given by Equation S3 and is larger when the values of the GLCM are concentrated in a few elements. The energy parameter is therefore an indicator of orderliness in the image.

$$\sum_{i,j=0}^{N-1} {P_{i,j}}^2 \tag{S3}$$

##### 4. Entropy

The quantity of homogeneity is given by Equation S4 and operates under the assumption that $\ln P_{i,j} = 0$. The entropy parameter is larger when there is greater disorder in the image.

$$\sum_{i,j=0}^{N-1} P_{i,j}(-\ln P_{i,j}) \tag{S4}$$

#### 5. Homogeneity

The quantity of homogeneity is given by Equation S5. The homogeneity parameter is larger when more elements are on the central diagonal due to the presence of the $(i-j)^2$ factor in the denominator, and is, therefore, an indicator of the image having similar pixel intensity values.

$$\sum_{i,j=0}^{N-1} \frac{P_{i,j}}{1+(i-j)^2} \tag{S5}$$

### S3. SEGMENTATION OF IMAGES FOR DATA ANALYSIS AND FOV SCORING

Due to the collagen imaged with SHG not always filling the entire image, SHG subimages found not to contain enough collagen were discarded through a manual review. An example is provided in Fig. S3(a), where the subimages designated A1, B1, B2, C1, and C2 are discarded due to insufficient collagen. The majority consensus for the later classification of the FOVs was determined by analyzing the four remaining subimages.

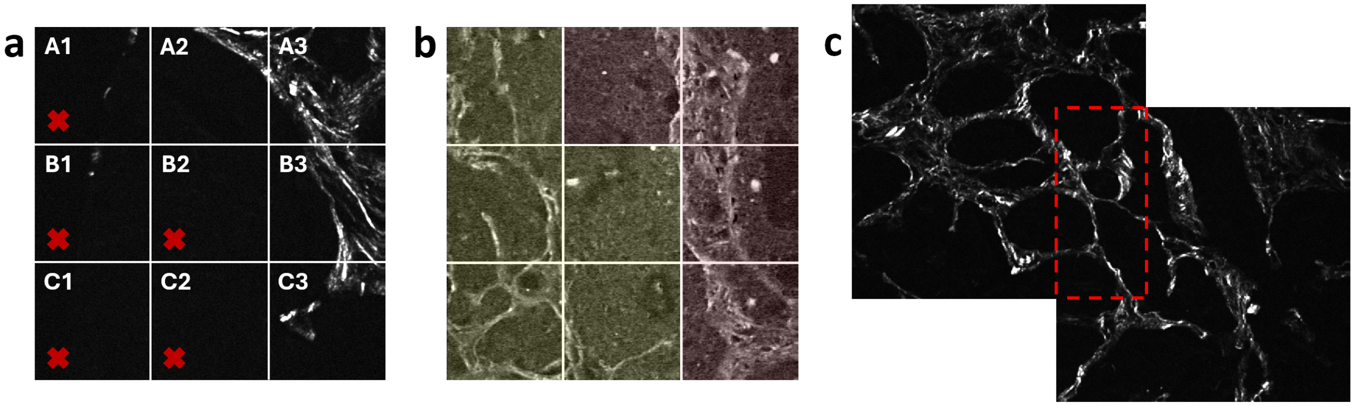


FIG. S3 SHG and SRS fields of view before segmentation. a) An SHG image is segmented into 3 x 3. Five of the nine subimages are discarded due to insufficient collagen. b) An image contains more than one tissue type. The image is split into HGC (yellow) and IDC (red) during segmentation. c) Two FOVs are overlapping. When overlapping images are segmented, the additional copies of the coincident subimages are discarded to avoid repeating data.

Fig. S3(b) is an example of an image where the field of view (FOV) was determined to contain more than one tissue type (in this case, HGC and IDC). In this case, 5 of the constituent subimages (when using the notational convention of Fig. S3(a): the subimages denoted A1, B1, B2, C1, and C2) were assigned a tissue type of HGC, and the remaining 4 subimages (the subimages denoted A2, A3, B3, and C3) were assigned a tissue type of IDC. The majority consensus for the later classification of the FOVs was then performed for two distinct FOVs, corresponding to the two groups of 5 and 4 subimages.

In Fig. S3(c), an example is shown where two FOVs overlap. This overlap would cause some of the subimages to be copies of the same region when segmented. To avoid this problem, additional copies of coinciding subimages are manually removed. Caution was taken to remove an equitable portion of subimages in each FOV such that subimages were not disproportionately removed from a single FOV. In the example shown in the Figure, when using the notational convention of Fig. S3(a), the subimage denoted C3 was removed from the upper left FOV, while the subimage denoted A1 was removed from the bottom right FOV.

### S4. KRUSKAL-WALLIS RANKED SUM TEST

The Kruskal-Wallis ranked sum test evaluates the p-value that two groups of statistics originate from the same distribution. $p$-values below 0.05 are considered significant. The tables in this section show, for each pairing of tissues (for example, IDC-P versus HGC) the $p$-value that the statistics derived from the subimages did not originate from a single distribution. For example, in the test performed on SHG subimages, a $p$-value of 0.04 was obtained for the contrast when comparing IDC-P and LGC images. This result indicates that the Kruskal-Wallis test evaluated a low probability that the contrast values from IDC-P and LGC subimages originate from the same distribution. Contrast can therefore be deduced to have statistical significance for distinguishing these two types of tissue.

TABLE S1 p-values obtained for SHG subimages with Kruskal-Wallis test. Values not meeting the threshold for statistical significance ($p < 0.05$) are shown in red.

| Statistic | p-values | | | | | |
|---|---|---|---|---|---|---|
| | IDC-P vs. HGC | IDC-P vs. LGC | IDC-P vs. Benign | HGC vs. LGC | HGC vs. Benign | LGC vs. Benign |
| Mean | 0.127 | 0.889 | $3.46\times10^{-4}$ | 0.331 | $9.97\times10^{-8}$ | $1.86\times10^{-5}$ |
| Variance | 0.985 | 0.878 | $1.08\times10^{-4}$ | 0.479 | $6.69\times10^{-6}$ | $3.23\times10^{-4}$ |
| Skewness | 0.496 | 0.682 | 0.989 | 0.997 | 0.965 | 0.984 |
| Kurtosis | 0.739 | 0.832 | 0.999 | 0.999 | 0.890 | 0.917 |
| Contrast | 0.999 | 0.040 | $3.90\times10^{-6}$ | 0.001 | $5.00\times10^{-7}$ | $3.65\times10^{-11}$ |
| Correlation | 0.991 | 0.487 | $1.37\times10^{-6}$ | 0.401 | $2.73\times10^{-7}$ | $3.19\times10^{-5}$ |
| Energy | 0.901 | $5.31\times10^{-11}$ | $4.04\times10^{-13}$ | $2.01\times10^{-16}$ | $3.68\times10^{-17}$ | $< 10^{-25}$ |
| Homogeneity | 0.067 | $4.64\times10^{-21}$ | $1.87\times10^{-10}$ | $1.87\times10^{-22}$ | $1.12\times10^{-17}$ | $< 10^{-25}$ |
| Entropy | 0.997 | $3.31\times10^{-4}$ | $1.79\times10^{-12}$ | $7.69\times10^{-8}$ | $2.62\times10^{-14}$ | $< 10^{-25}$ |

TABLE S2 p-values obtained for 1450 $cm^{-1}$ subimages with Kruskal-Wallis test. Values not meeting the threshold for statistical significance ($p < 0.05$) are shown in red.

| Statistic | p-values | | | | | |
|---|---|---|---|---|---|---|
| | IDC-P vs. HGC | IDC-P vs. LGC | IDC-P vs. Benign | HGC vs. LGC | HGC vs. Benign | LGC vs. Benign |
| Mean | $1.32\times10^{-4}$ | $5.26\times10^{-5}$ | 0.814 | 0.693 | 0.662 | 0.366 |
| Variance | 0.275 | 0.002 | 0.952 | 0.092 | 0.988 | 0.467 |
| Skewness | 0.647 | 0.203 | 0.374 | 0.663 | 0.113 | 0.037 |
| Kurtosis | 0.182 | 0.003 | 0.155 | 0.182 | 0.009 | $3.93\times10^{-4}$ |
| Contrast | 0.073 | $4.46\times10^{-8}$ | $1.33\times10^{-7}$ | $9.26\times10^{-5}$ | $2.12\times10^{-5}$ | 0.106 |
| Correlation | 0.134 | $2.99\times10^{-8}$ | 0.888 | $2.06\times10^{-5}$ | 0.991 | 0.050 |
| Energy | 0.005 | $1.48\times10^{-11}$ | 0.042 | $6.10\times10^{-6}$ | 0.666 | 0.546 |
| Homogeneity | 0.100 | $1.29\times10^{-4}$ | 0.001 | 0.036 | 0.029 | 0.566 |
| Entropy | 0.002 | $1.34\times10^{-17}$ | 0.016 | $3.31\times10^{-10}$ | 0.514 | 0.220 |

TABLE S3 p-values obtained for 1668 $cm^{-1}$ subimages with Kruskal-Wallis test. Values not meeting the threshold for statistical significance ($p < 0.05$) are shown in red.

| Statistic | p-values | | | | | |
|---|---|---|---|---|---|---|
| | IDC-P vs. HGC | IDC-P vs. LGC | IDC-P vs. Benign | HGC vs. LGC | HGC vs. Benign | LGC vs. Benign |
| Mean | $4.28\times10^{-9}$ | $5.56\times10^{-8}$ | 0.444 | 0.865 | $3.25\times10^{-5}$ | $1.63\times10^{-5}$ |
| Variance | $1.36\times10^{-4}$ | $1.65\times10^{-6}$ | 0.351 | 0.230 | $9.28\times10^{-4}$ | $3.88\times10^{-5}$ |
| Skewness | $4.48\times10^{-5}$ | 0.079 | 0.204 | 0.457 | 0.999 | 0.936 |
| Kurtosis | $9.95\times10^{-6}$ | 0.986 | 0.027 | $2.45\times10^{-4}$ | 0.939 | 0.051 |
| Contrast | $< 10^{-25}$ | $< 10^{-25}$ | 0.158 | $2.10\times10^{-6}$ | $< 10^{-25}$ | $3.79\times10^{-16}$ |
| Correlation | $5.34\times10^{-5}$ | $8.72\times10^{-11}$ | 0.992 | 0.002 | 0.244 | 0.003 |
| Energy | $1.48\times10^{-12}$ | 0.787 | 0.327 | $4.63\times10^{-8}$ | $6.43\times10^{-7}$ | 0.125 |
| Homogeneity | $< 10^{-25}$ | $3.49\times10^{-19}$ | 0.240 | 0.001 | $2.32\times10^{-20}$ | $2.96\times10^{-11}$ |
| Entropy | $2.79\times10^{-18}$ | 0.963 | 0.074 | $6.76\times10^{-14}$ | $6.22\times10^{-11}$ | 0.040 |

### S5. ACCURACY OF SVM MODELS WHEN USING EITHER FIRST-ORDER OR SECOND-ORDER PARAMETERS ONLY

In the feature selection portion of the paper, models were trained with all parameters except one present to evaluate the importance of these parameters in the SVMs. A similar approach can be applied to determine the relative importance of the first-order parameters (mean, variance, skewness, kurtosis) and second-order parameters (contrast, homogeneity, energy, correlation, and entropy) to the classification accuracies of the SVM models. Table S4 shows the mean classification accuracies of the SVM models trained on SHG, 1450 $cm^{-1}$ SRS, and 1668 $cm^{-1}$ SRS subimages when training on either first-order or second-order parameters only.

TABLE S4 Mean accuracy results for SVM models trained on SHG, 1450 $cm^{-1}$ SRS, and 1668 $cm^{-1}$ SRS subimages when training on either first-order or second-order parameters only. The mean classification accuracy of the central diagonal of the confusion matrix representing subimage accuracy as a function of which parameters are trained on. Mean accuracies are shown in contrast to the accuracy when incorporating both first-order and second-order statistics, shown in bold.

| Statistics in SVM model | Overall classification accuracy (%) | | |
|---|---|---|---|
| | SHG SVM model | 1450 $cm^{-1}$ SRS SVM model | 1668 $cm^{-1}$ SRS SVM model |
| **All parameters** | **92.60** | **49.30** | **85.40** |
| First-order parameters only | 32.15 | 24.18 | 29.34 |
| Second-order parameters only | 90.09 | 27.78 | 40.04 |

When training the SVM models on only the first-order statistics, the mean classification accuracy for the SVM model trained on SHG subimages is 32.2%, the mean classification accuracy for the SVM model trained on 1450 $cm^{-1}$ SRS subimages is 24.2%, and the mean classification accuracy for the SVM model trained on 1668 $cm^{-1}$ SRS subimages is 29.3%. These results are scarcely better than the 25% that would be expected from random chance, indicating that gray-level statistics do not possess much discriminatory power for the different tissue types when used without texture analysis.

When training the SVM models on only the second-order statistics, the mean classification accuracy for the SVM model trained on SHG subimages is 90.1%, the mean classification accuracy for the SVM model trained on 1450 $cm^{-1}$ SRS subimages is 27.8%, and the mean classification accuracy for the SVM model trained on 1668 $cm^{-1}$ SRS subimages is 40%. For the SVM model trained on SHG subimages, the texture parameters alone allow the model to reach a mean accuracy that is only 2.5% lower than the model using all parameters, indicating that texture does the bulk of tissue discrimination. In the case of the SVM model trained on 1450 $cm^{-1}$ SRS subimages, the classification accuracy remains in the ballpark of random chance, reflecting the poor overall classification accuracy of the 1450 $cm^{-1}$ SRS subimages to begin with. In the case of the SVM model trained on 1668 $cm^{-1}$ SRS subimages, using only texture parameters provides a mean accuracy that is clearly superior that achieved using only first-order parameters, but that is still less than half of the mean accuracy when all parameters are used. It can be concluded that, in the case of the 1668 $cm^{-1}$ SRS subimages, the SVM model relies on a mixture of both first-order and second-order parameters to achieve its high accuracies.
